# Regulation of CtISWI activity by AutoN and HSS domain

**DOI:** 10.64898/2026.01.25.701572

**Authors:** Yiming Zhao, Wansen Tan, Jingjun Hong

## Abstract

Chromatin remodeler imitation switch (ISWI) plays an important role in regulating chromatin structure through sliding and spacing nucleosomes. Despite the enormous progress in regulatory elements and mechanisms of the activity of ISWI in recent years, there are still some unclear structures and mechanisms in different species. Here, we studied the ATPase activity and nucleosome binding affinity of *Chaetomium thermophilum* ISWI (hereafter referred to as *Ct*ISWI^WT^) and several mutants, further proving the importance of these mutated residues in the inhibition of AutoN. We also analyzed the effects of dsDNA and ssDNA on ATPase activity of *Ct*ISWI, suggesting the potential interaction between HSS and ATPase domain. Notably, we provided a predicted structural model based on the sequence of *Ct*ISWI^WT^, proposing a two-step activating mechanism of conformation change and activity regulation. Taken together, our findings elucidate a different model of ISWI self-maintenance and action, providing a new mechanism of regulation supporting chromatin remodeling.

**Highlights:**

1. Structural modeling of ISWI: a chromatin remodeler that couples to ATP hydrolysis to slide and space composition of nucleosome.
2. The ATPase activity of ISWI can be stimulated by exogenous DNA, with opposite promoting effects by dsDNA and ssDNA.
3. The mechanism by which ISWI is activated upon binding with nucleosome has been the subject of debate, and a more comprehensive mechanism for regulation of ISWI activity.

**Significance:** In the past decades, a variety of regulatory mechanisms of the activity of chromatin remodeling factor ISWI have been proposed. Based on the hypothesis of nucleosome complex structure analysis, these studies attempted to explore the mechanism of chromatin remodeling, a gene expression regulation activity. However, previous studies have basically focused on the binding and regulation mechanism of ISWI ATPase domain and nucleosomes, without mentioning the activity mode of full-length ISWI. Therefore, our study mainly focuses on the nearly full-length ISWI containing HSS domain, exploring the mechanism of the active state transition of ISWI in remodeling activities from this perspective. It enriches and supplements the research on chromatin remodeling, an important physiological activity.

## 1. Introduction

ISWI plays a role in gene regulation processes such as DNA replication and transcription, heterochromatin formation, and nucleosome remodeling. The ATPase domain occupies the main body of the entire subunit and plays a major catalytic role in remodeling reactions. ISWI mainly achieves chromatin remodeling by consuming the energy generated by ATP hydrolysis to push nucleosomes to slide, finally arranging nucleosomes at hierarchical intervals on DNA. Structurally, the ATPase domain consists of two parts, core 1 and core 2, which can independently bind to nucleosomes and are not affected by other domains. After binding, it causes a conformational change in itself, thereby activating ATP hydrolysis ^[1]^. Moreover, the energy released from ATP hydrolysis is not only used to promote nucleosome sliding, but also to drive “navigation” movement of ISWI in compacted chromatin. ISWI achieves movement by continuously alternating binding and release with multiple sites on different nucleosomes in the array. This directly confirms its ability to stably bind to nucleosomes in a condensed structure ^[2]^.

There are two self-inhibitory regions, AutoN and NegC, distributed on both sides of ATPase, which together with the ATPase domain form the catalytic core of ISWI. It was found that the sequence of AutoN is similar to the basic patch of H4, and its inhibitory effect can be relieved by the basic patch ^[3]^. It was found that the inhibitory effect of AutoN is reflected in hindering ATP hydrolysis, and NegC inhibits nucleosome sliding by uncoupling ATP hydrolysis and DNA translocation. However, inconsistent with this binding mode, after *Ct*ISWI_77-Δ-722_ binds to nucleosomes, the structure of AutoN become disordered. Due to the binding of nucleosome, the structure of AutoN undergoes significant changes ^[4]^. After *Saccharomyces cerevisiae* ISW1 binds to nucleosomes, conformational changes also occur. Under the induction of ISW1, the H4 tail of nucleosome will swing to the vicinity of the negatively charged surface of core 2 and bind to it ^[5]^. The sensitivity of ISWI to flanking DNA is closely related to the ability of nucleosomes to be moved to a certain position ^[6,7]^, and NegC also plays an important role in regulating the sensitivity of ISWI to linker DNA. When NegC of SNF2h is replaced by an equal length linker, the overall remodeling activity of SNF2h lacking NegC is not significantly affected, but its sensitivity to Flanking DNA is severely reduced. Compared with the wild-type SNF2h, the speed difference of this mutated SNF2h reconstruction of nucleosomes without linker DNA and 30/30 nucleosomes is smaller ^[8]^. During ATP hydrolysis and remodeling, NegC promoted a pause state of ISWI action in the remodeling cycle ^[9]^. Consistent with it, residues 643-648 of NegC had high flexibility in ISWI in the absence of cofactors, which is at the same position in CHD1, and their deletion only slightly reduced the inhibitory effect ^[10]^.

There are three nucleosome binding sites at the C-terminus of ISWI, namely HAND, SANT, and SLIDE, which together form a dumbbell shaped HSS domain ^[11]^. The main function of HSS is to bind nucleosome linker DNA, enhance the affinity and reaction rate of ISWI, assist in sensing flanking DNA, and promote specific effects in the reaction ^[1]^. During chromatin remodeling, ISWI often forms remodeling complexes with different accessory subunits. These accessory subunits can provide additional functions for the complex, helping to recognize specific substrates and produce different catalytic effects, thereby improving the specificity of the reaction ^[12]^. In addition, accessory subunits also have a certain regulatory effect by binding to nucleosome epitopes and jointly regulating catalytic processes with other components ^[13]^.

Although the components above can independently affect chromatin remodeling, various factors often constrain each other and play a comprehensive role in the remodeling response.

Flanking DNA usually uses its length as a scale for the transformation of each regulatory link. The key regulatory epitopes for the activity of ISWI are linker DNA and H4 tail of nucleosome. While AutoN inhibits ISWI activity, its own function is also controlled by H4 tail and HSS ^[3]^. In ACF, H4 competes with AutoN for binding to Snf2h and Acf1 subunits. When the DNA length is insufficient, Acf1 preferentially binds to H4 tail, and AutoN can therefore bind to SNF2h, inhibiting ATPase activity ^[14]^. When the DNA is long enough, HSS binds to linker DNA, which not only promotes the binding of lobe 2 to H4 and nucleosome DNA, but also breaks down the structure of NegC, releasing inhibition, and promotes the activity of ATPase domain. Due to conformational changes in SNF2h, DNA moves around the nucleosome, linker DNA shortens, causing HSS to detach, restoring the structure of NegC and allowing it to bind to lobe 2, inducing the formation of an inactive conformation ^[8]^.

Here we mutated some key residues based on the sequence similarity (Supplementary data Figure S3) between *Ct*ISWI and *Mt*ISWI, combined with original existing interaction between AutoN and core 2 (Figure 2A), then measured the ATPase activity of the wild-type and each mutant. We also used exogenous dsDNA and ssDNA to stimulate protein activity and compared it before and after stimulation. Besides, nucleosome-binding affinity of these mutants was also tested to prove the importance of those residues. What’s more, based on our predicted structure of *Ct*ISWI^WT^, we proposed a new mechanism for the autonomous regulation of *Ct*ISWI activity, combined with previous studies.

## 2. Materials and methods

### 2.1 Protein expression and purification

*Ct*ISWI_77-1038_ (construct 1 in Figure 1A) and the catalytic core of *Ct*ISWI_77-722_ (*Ct*ATPase, construct 2 in Figure 1A) were used in most of the biochemical analyses (Supplementary data Figure S1). To generate the 8G/R140A, 8G/3A and 8G/4A mutants, we firstly replaced the residues 133-136 and 148-151 of AutoN with two sequences of GGGG respectively, then generated R140/A, and E155 - D157/A based on 8G mutant. Various point mutants were generated by PCR (Supplementary data Table S1-4). All constructs were confirmed by DNA sequencing. Each protein was purified by Ni-affinity chromatography and FPLC (Supplementary data Figure S2). More detailed steps and content of the protein expression and purification have been mentioned in the protocol we previously published ^[15]^.

**Figure 1.**
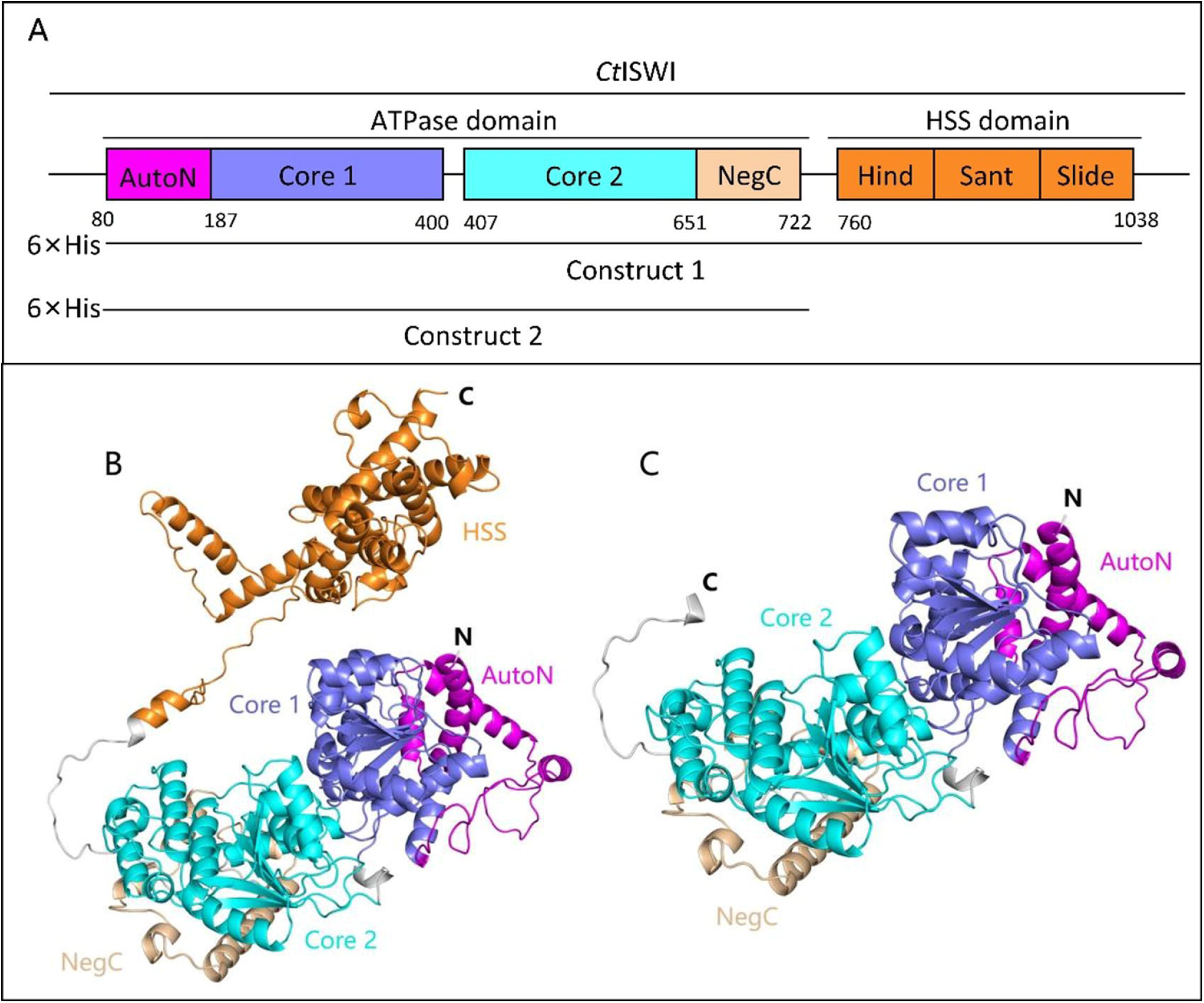
Overall structure model of *Ct*ISWI and *Ct*ATPase. **(A)** Domain architecture of *Ct*ISWI. **(B)** Predicted structural model of *Ct*ISWI_77-1038_; **(C)** Predicted structural model of *Ct*ATPase_77-722_.

### 2.2 ATPase activity assays

Measurement of ATP hydrolysis was based on a spectrophotometric shift in the absorbance of the substrate under 660 nm, resulting from the production of inorganic phosphorus by ATP hydrolysis in the presence of Ca^2+^ Mg^2+^ ATPase (Ca^2+^ Mg^2+^ ATPase activity assay kit). The measurements were performed in an ultraviolet spectrophotometer, and the ATPase activities were calculated at the early time points when the yield of product increased linearly. ATPase assays were performed in 150 mM NaCl, 10 mM Tris-HCl (pH 7.2), and 1 mM MgCl_2_. Owing to the auto-inhibited nature and the low activities, 10 μM of *Ct*ATPase_77–722_ and various *Ct*ISWI_77– 1038_ proteins, including the enzymes with wild-type interface and with various mutations, were used. Owing to high ATPase activities caused by the promotion of exogenous dsDNA or ssDNA, 10 μM *Ct*ISWI and 1 μM DNA were used. The specific ATPase activities of all the proteins were normalized to *Ct*ISWI without DNA. More details and content of measurement of ATPase activity have been mentioned in the protocol we previously published ^[15]^.

### 2.3 Mononucleosome preparation and purification

Monoucleosome with 167 bp DNA was used in binding assays, with 10 bp linker DNA on both sides of nucleosome bound by HSS domain of ISWI. More details and content of histone octamer purification and mononucleosome preparation have been mentioned in the protocol we previously published ^[16]^.

### 2.4 Nucleosome binding assays

Nucleosome binding assays were performed in the buffer (10 mM Tris-HCl, pH 7.2, 2 mM β-ME, 1 mM MgCl_2_), and *Ct*ISWI mutants were on dialysis against 75 mM NaCl, 10 mM Tris-HCl (pH 7.2) before assays. The final concentration of nucleosomes used was controlled at 0.1 μM. The reactions were performed on ice with 0.1 μM nucleosomes and *Ct*ISWI with concentration gradients. The products were resolved with 6% native acrylamide gels, 0.2×TBE at 4°C for 150 min at 90 V. The positions of fluorescently labelled DNA were detected using a gel imaging analysis system (Miulab).

### 2.5 Protein structural model prediction

The structural model of *Ct*ISWI_77-1038_ and *Ct*ATPase_77-722_ was predicted and assessed by the SWISS-MODEL server (https://swissmodel.expasy.org/assess/9hV9b9/01). SWISS-MODEL predicted a total of 6 models, but we selected 01 model, named model 1, because our experiment matches this model well. The structure model of *Ct*ISWI (UniProt ID: G0S9L5) predicted by AlphaFold in UniProt database was named model 2 (AF-G0S9L5-F1).

## 3. Results

### 3.1 Relative ATPase activity of *Ct*ISWI^WT^ and mutants *in vitro*

Studies in thermophilic yeast, the wild type *Mt*ISWI_81–1048_ has a very poor ATPase activity *in vitro*, repressed by AutoN through multiple mechanisms ^[17]^. Based on a higher proportion of amino acid sequence homology, the ATPase activity of *Ct*ISWI_77-1038_ our study (Figure 1) is also very poor. In order to alleviate the inhibitory effect of AutoN on protein activity as much as possible, based on our own sequence (Figure 2A), a series of mutants were designed to mutate some consecutive basic residues among AutoN into uncharged amino acids such as alanine or glycine (K_133-136_/G, R_148-151_/G, abbreviated as 4R/4G and 8G below). It is found that mutations of these residues increase the ATPase activity, although there is no significant difference between 4R/4G and 4K/4G alone (Figure 2D). To further enhance the activity of the protein, we chose some polar amino acid residues to mutate according to the key residues in *Mt*ISWI inhibition by AutoN (R_140_/A, and E_155_-D_157_/A abbreviated as 3A below) (Figure 2B). The result shows that ATPase activity of these mutants is significantly higher than the first three proteins (Figure 2D) (Supplementary data Table S7). These results suggest that K_133-136_, R_148-151_ are important for *Ct*ISWI inhibition, and R_140_, E_155_-D_157_ play an important role in the interaction between AutoN and core 2.

**Figure 2.**
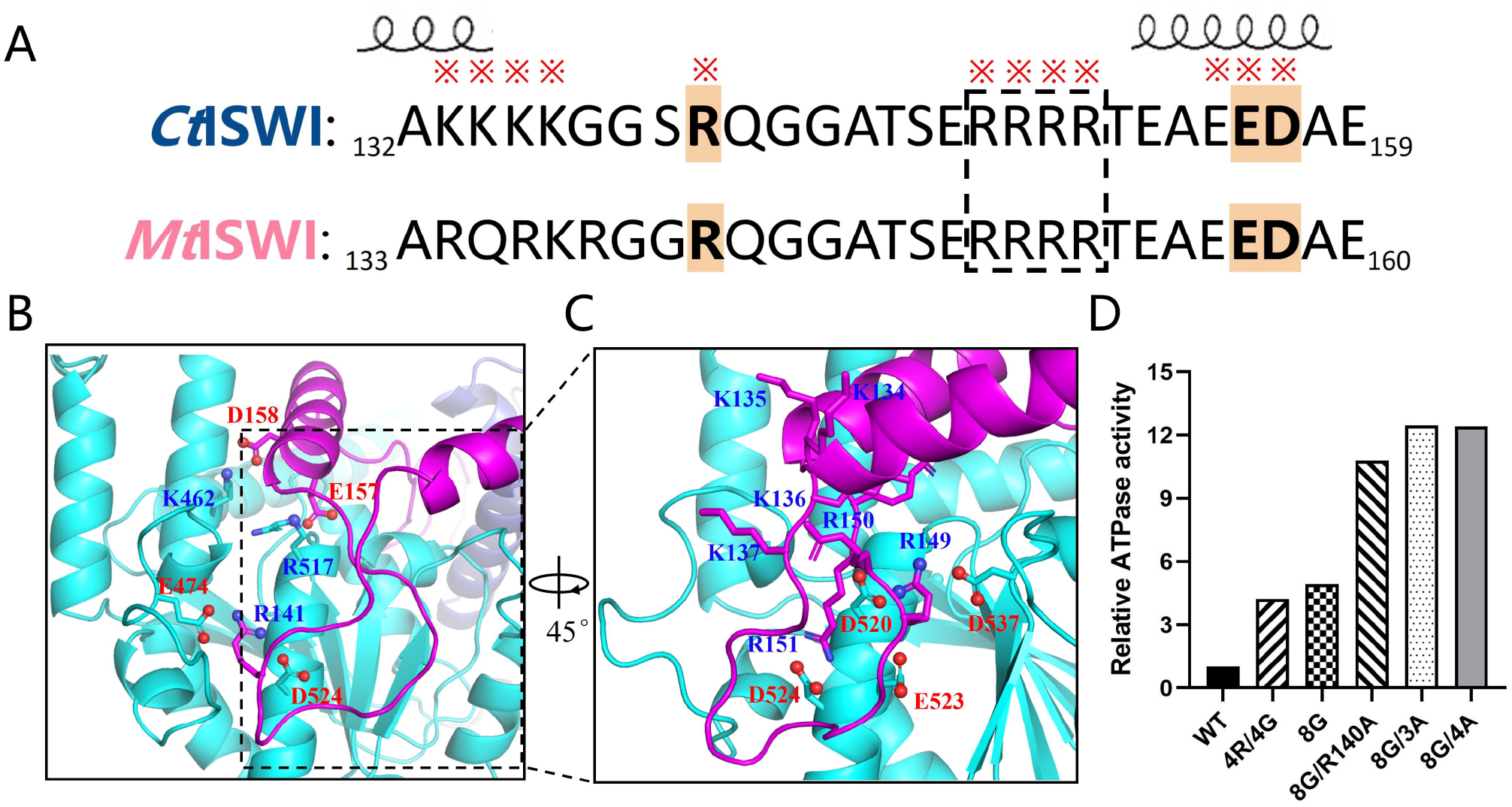
Mechanism of AutoN inhibition on ISWI. **(A)** Sequence alignment between *Ct*ISWI and *Mt*ISWI. The conserved residues involved in the interaction between AutoN and core 2 are highlighted. The residues mutated of *Ct*ISWI are highlighted by red star on the top. Mutated R148-151 region is marked in red frame. **(B)** Interaction between AutoN and core 2 domain of *Mt*ISWI (PDB: 5JXR). Based on the sequence alignment, R141, E157 and D158 correspond to R140, E156 and D157 mutated in our *Ct*ISWI. **(C)** Key residues among basic patch on AutoN and acidic patch on core 2 of *Ct*ISWI. R134-R136 of *Mt*ISWI have been mutated to K in this structure. **(D)** Relative ATPase activities of wild-type *Ct*ISWI and various mutants in AutoN. The specific activities of the proteins were normalized to wild type.

### 3.2 Promoted ATPase activity of *Ct*ISWI by exogenous DNA

To analyze the facilitation of DNA to ATPase activity of *Ct*ISWI, we first added extra 32 bp dsDNA and *Ct*ISWI according to a certain concentration ratio, when measuring its ATPase activity. Result shows that 32 bp dsDNA has an activating effect on the ATPase activity. Subsequently, we chose other dsDNA of different length (Supplementary data Table S5). It is found that dsDNA longer than 32 bp can also promote the ATPase activity and it seems to have a certain dependence on the length of dsDNA (Figure 3A, Supplementary data Table S8), whereas the shorter dsDNA has no significant effect. In addition to dsDNA, we explored whether ssDNA had an effect on ATPase activity of *Ct*ISWI (Supplementary data Table S6). We also put 33 nt ssDNA and protein together firstly, and we found that 33 nt ssDNA did promote the ATPase activity (Figure 3B, Supplementary data Table S9), then the other longer ssDNA was tested. Interestingly, although they have activating effect, the promotion is lower when the ssDNA is longer, which is contrary to dsDNA. Actually, research on S. cerevisiae (yIsw2) showed that yISW2 is an ATP-dependent single stranded DNA translocase ^[18]^. The max velocity of ssDNA stimulated ATP hydrolysis of Isw2 will increase with increasing DNA length, whereas the promotion effect gradually weakens ^[19]^.

**Figure 3.**
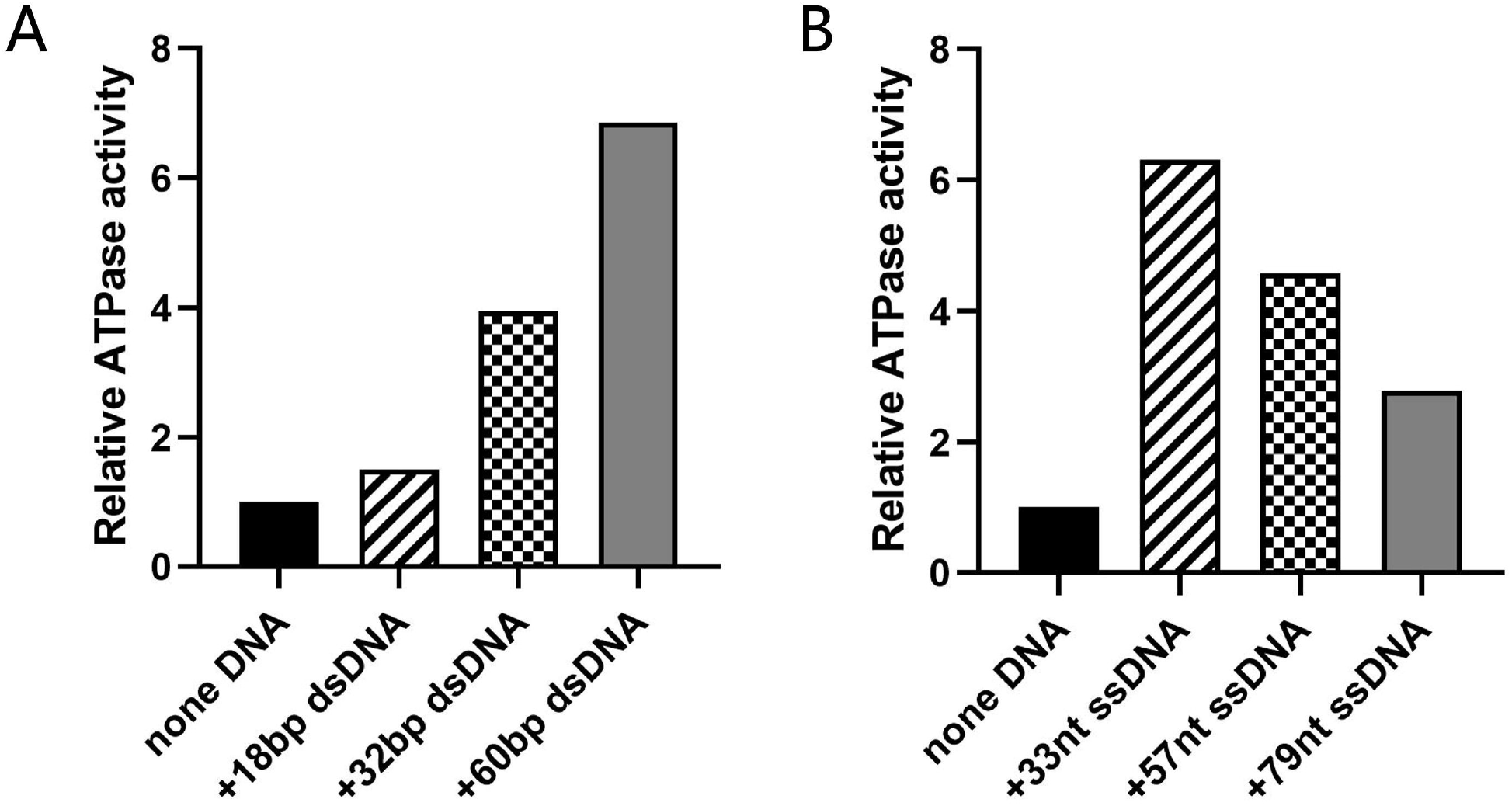
Promotion of exogenous DNA on ATPase activity of *Ct*ISWI. **(A)** Promotion of dsDNA with different length on *Ct*ISWI. **(B)** Promotion of ssDNA with different length on *Ct*ISWI. The specific activities of the proteins were normalized to the protein with no DNA.

### 3.3 Relative ATPase activity of the catalytic core *Ct*ISWI *in vitro*

Actually, to find out whether the ATPase domain has relatively high activity, we made a truncation at Ala723 of *Ct*ISWI_77-1038_ to get *Ct*ATPase_77-722_, which is the catalytic core of *Ct*ISWI. Surprisingly, the result shows that the activity of *Ct*ATPase_77-722_ is very low (Figure 4). There is still a significant difference between the activity of the ATPase domain and *Ct*ISWI, although the full length *Ct*ISWI also has a low activity which is shown above. From the structural perspective, ISWI has an additional HSS region at the C-terminus compared to ATPase. The HSS domain is indeed not necessary for the catalytic function of ISWI ^[1]^, which can assist in protein sensing and binding to flanking DNA, thereby increasing the affinity and reaction rate of ISWI, and promoting specific remodeling effects. This higher ATPase activity of *Ct*ISWI also suggests that the HSS domain does have an important stimulated impact on the activity of *Ct*ISWI. The binding function is mainly accomplished by SLIDE, which binds to DNA by forming a positively charged channel, helping ISW2 to accurately locate on nucleosomes ^[20]^. After binding to linker DNA, the SLIDE domain promotes the hydrolysis of ATP by catalytic subunits and pushes DNA into nucleosomes from the entrance, enhancing nucleosome translocation ability ^[21]^.

**Figure 4.**
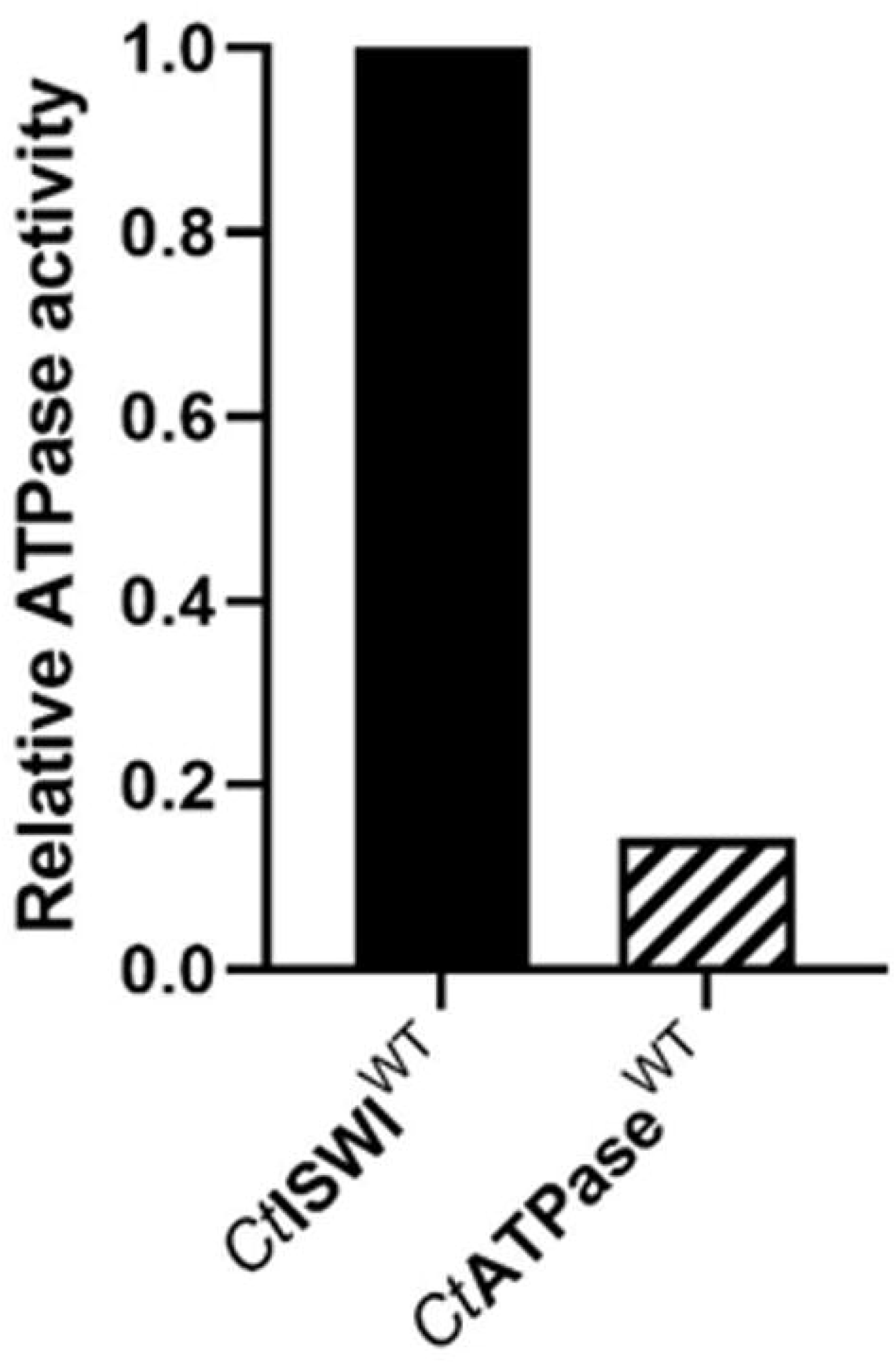
Relative ATPase activity between *Ct*ISWI and its catalytic core. The specific activities of the proteins were normalized to wild type.

### 3.4 Nucleosome binding activity of *Ct*ISWI *in vitro*

Besides, in order to analyze the function of *Ct*ISWI, nucleosome sliding assay was carried out. It was found out that wild-type *Ct*ISWI has almost no complex with nucleosome (Figure 5A). In sharp contrast, disruption of the salt-bridge interactions between AutoN and core 2 by R140A and (E155-D157)/A mutations (8G/R140A, 8G/3A and 8G/4A) resulted in stronger binding effect (Figure 5B-D), which is in consistent with the change in ATPase activity (Figure 2D). These results further suggest that the AutoN-core 2 interactions are essential for overall activity of *Ct*ISWI.

**Figure 5.**
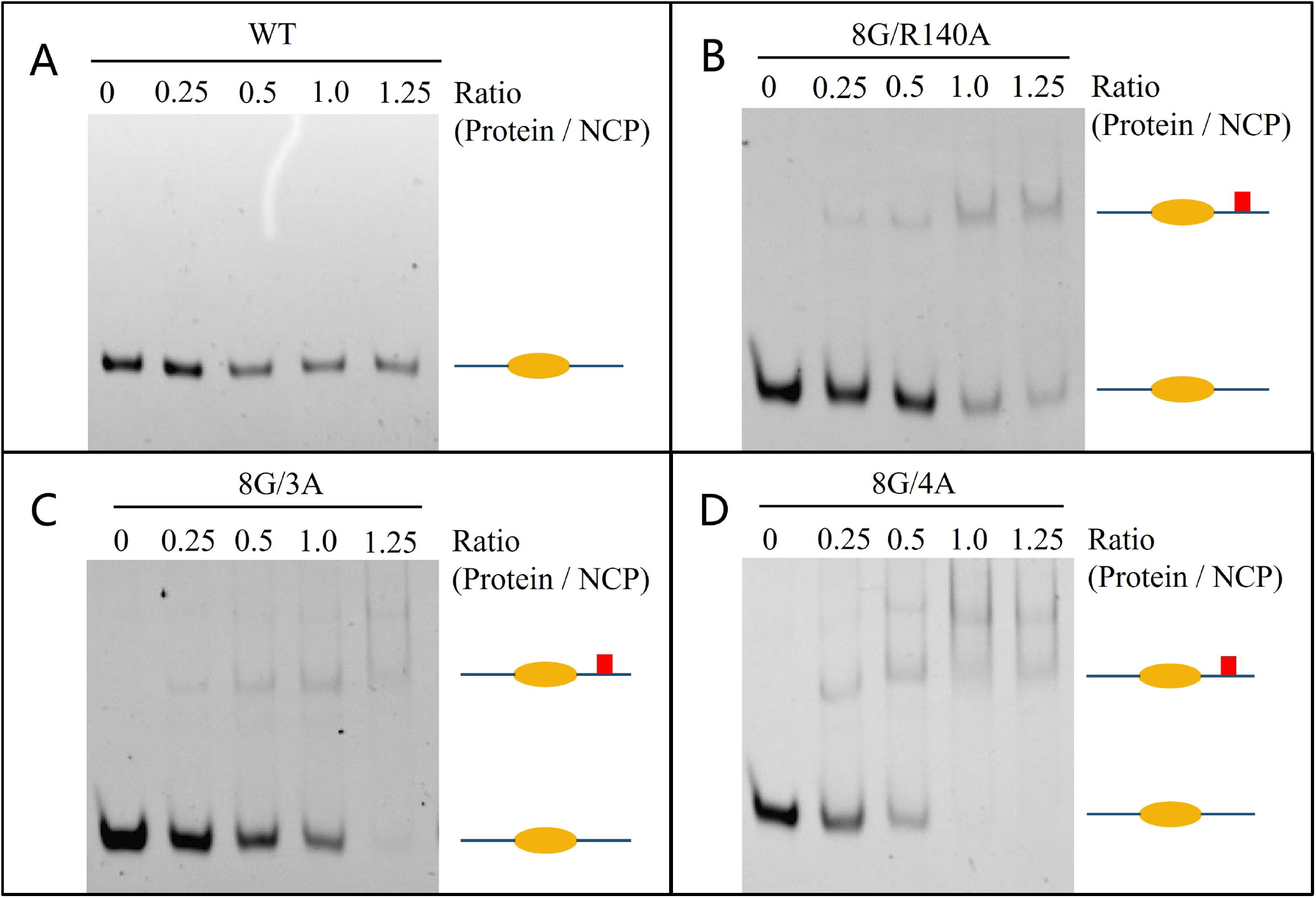
Nucleosome binding activity of *Ct*ISWI wild type and mutants. **(A)** Gels of the nucleosome-binding assays of wild type *Ct*ISWI. **(B-D)** Gels of the nucleosome-binding assays of three mutants. Three sets of independent experiments were performed, and the representative one is shown.

## 4. Discussion

Taken together, our study revealed the inhibition of AutoN on the ATPase activity of *Ct*ISWI once again. Through the preparation of corresponding mutants and their activity measurement, we found that these residues play an important role in ATP hydrolysis and the binding with nucleosome. Our findings also suggested the promotion of exogenous dsDNA to ATPase activity of *Ct*ISWI. Previous study has revealed that linker DNA between two nucleosomes can stimulate the activity of ISWI, as DNA becomes longer, the rate at which ISWI catalyzes ATP hydrolysis and DNA translocation increases ^[22]^. In the absence of nucleosome, we chose exogenous dsDNA to simulate linker DNA of nucleosome. ATPase activity of *Ct*ISWI can be promoted by dsDNA with a certain length (longer than about 30 bp) ^[23]^. In chromatin background, the chromatin constructed by 25 bp DNA connection is more prone to stable phase separation, has strong viscoelasticity, and is highly similar to the high-density region of natural chromatin. On the contrary, the overall network of 30-bp system is looser, exhibiting higher dynamism, faster molecular diffusion, and easier modification and disintegration of histones ^[24]^. Moreover, the absence of H4 tail has a much greater impact on the 30 bp fiber structure than on the 25 bp. These findings also confirm that chromatin has a more open and active conformation, which facilitates remodelers such as ISWI to perform higher remodeling functions when linker DNA is longer than 30 bp. Cells can also switch the physical state of the genome by adjusting the distance between nucleosomes, which affects gene expression and genetic stability.

In previous study, the structure of *Ct*ISWI_77-Δ-722_ binding with nucleosome (6PWF) is different from *Mt*ISWI (5JXR). Upon fixing the position of core 1, core 2 has a 137° deflection after binding with nucleosome ^[4]^ (Supplementary data Figure S5A-B). Here, our model of *Ct*ISWI predicted by SWISS-MODEL (model 1) and that by AlphaFold (model 2) are consistent with *Ct*ISWI_77-Δ-722_ (6PWF) (Supplementary data Figure S5A, S5C, S5D). There are differences in the position of HSS domain in model 1 and model 2, which may represent two different states of *Ct*ISWI (Supplementary data Figure S5C-D). Our data shows that there may be interactions between HSS domain and core 1 or core 2 (Figure 4). Measuring the distance between HSS domain and core 1, it is found that HSS seems to be closer to core 1 in model 1 than model 2 (Supplementary data Figure S5E-F), which makes us trust model 1more than model 2. What’s more, based on the superposition of *Ct*ISWI model with nucleosome (6PWF), it is found that there are space conflicts between the nucleosome and HSS domain (Supplementary data Figure S5G-H), which may explain why our 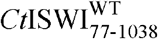 can hardly bind with nucleosome.

To further understand the relationship between HSS domain and *Ct*ATPase, we presented the surfaces of model 1 and model 2. In model 1, the HSS domain is biased towards Core 1 and located above it (Supplementary data Figure S6A), while in model 2, the HSS domain is located above the junction of Core 1 and Core 2, compared to Core 2, the HSS domain is closer to Core 1 (Supplementary data Figure S6B). Comparing the HSS domains of model 1 and model 2, it is evident that they bind to different positions of *Ct*ATPase, indicating a conformational change. The movement of HSS domain is the structural basis regulated by Linker DNA when ISWI binds to nucleosomes (Supplementary data Figure S6).

In addition, our study also suggested that ssDNA has a promoting effect on the ATPase activity of *Ct*ISWI. Surprisingly, the trend of this function is opposite to dsDNA, that is, as ssDNA prolongs, the promoting effect weakens. The length of nucleosome linker DNA in eukaryotes is highly variable, typically ranging from 10-80 bp, with an average of approximately 33 bp ^[25]^. When the DNA is shorter than 80 bp, ISWI seems to receive a kind of “instruction” to increase its activity and move nucleosome to an optimal position. Whereas, as the length of DNA increases, the need for nucleosome movement decreases, so the promoting effect of DNA on the activity of ISWI will also weaken. By the way, it is speculated that the effect of ssDNA on ATPase activity of *Ct*ISWI might stem from the binding of ISWI with ssDNA that briefly exists after unwinding during DNA transcription. Our results demonstrated the regulatory role of HSS domain in ISWI activity by recognizing DNA once again, regardless of the stimulating effect of dsDNA or ssDNA on ISWI activity.

A crosstalk phenomenon between Snf2H oligomers has been previously observed in the cryo-EM structure of double bound nucleosomes [^26, 27^]. This asymmetric binding of two ISWI molecules seems to be an inherent characteristic of nucleosome interactions. Our data showed that for *Ct*ISWI^8G/3A^ and *Ct*ISWI^8G/4A^, as the ratio of protein is higher, the nucleosome might tend to bind to two *Ct*ISWI (Figure 5C-D). In previous studies, the ISWI construct was added to the nucleosome in a 2:1 ratio ^[28]^. Two ATPase motors were bound to the opposite ends of the nucleosome (SHL+2/-2) then moved in opposite directions, establishing an effective dynamic equilibrium between the ground state prior to activation and translocation state of histone octamers.

Although we were unable to get the complex structure of *Ct*ISWI and nucleosomes, based on structural prediction model, we found that the HSS domain of *Ct*ISWI will fold back physically to the ATPase domain. That’s why our *Ct*ISWI still has a low affinity for nucleosomes, even after the corresponding residue mutation of AutoN.

Previous study has illustrated nucleosome recognition by the ATPase core domain and linker DNA binding by HSS domain ^[28]^. ISWI is an intrinsically dynamic protein during the catalytic process, relying on conformational changes including AutoN, core 1 and core 2, NegC and HSS regions ^[30]^. When there are no nucleosomes or nucleosome DNA with different lengths, ISWI exhibits various dynamic structural changes. ATPase core domains are sufficient to induce conformational changes, and the degree of these changes depends on the presence of NegC and nucleotide states ^[10]^. Besides, there is also a significant loss of histone signal intensity when SNF2h binds in presence of ADP-BeFx, indicating that histone octamers have significant conformational dynamics and plasticity ^[28]^. Although the explanation of the functions of each region in ISWI is clear, there is not enough description about the dynamic progress of conformational change and remodeling. Thus, a description of the detail during the progress of ISWI conformational change is necessary.

Based on our predicted model, we propose a regulatory mechanism for the transition of ISWI from ground state to activated state, and nucleosome is the key element in this regulatory process. Free ISWI adopts a highly dynamic, auto-inhibited conformation (Figure 6A a, d), representing an inactive state. In one conformation, HSS domain of ISWI covers the cleft between core 1 and core 2, preventing nucleosomes from approaching ISWI. In another conformation, AutoN crosses the cleft between core 1 and core 2 with a triangular shape, locking them together and inhibiting the activity of core 1, thereby preventing efficient ATP hydrolysis. However, the order in which these two auto-inhibited conformations appear needs further study.

**Figure 6.**
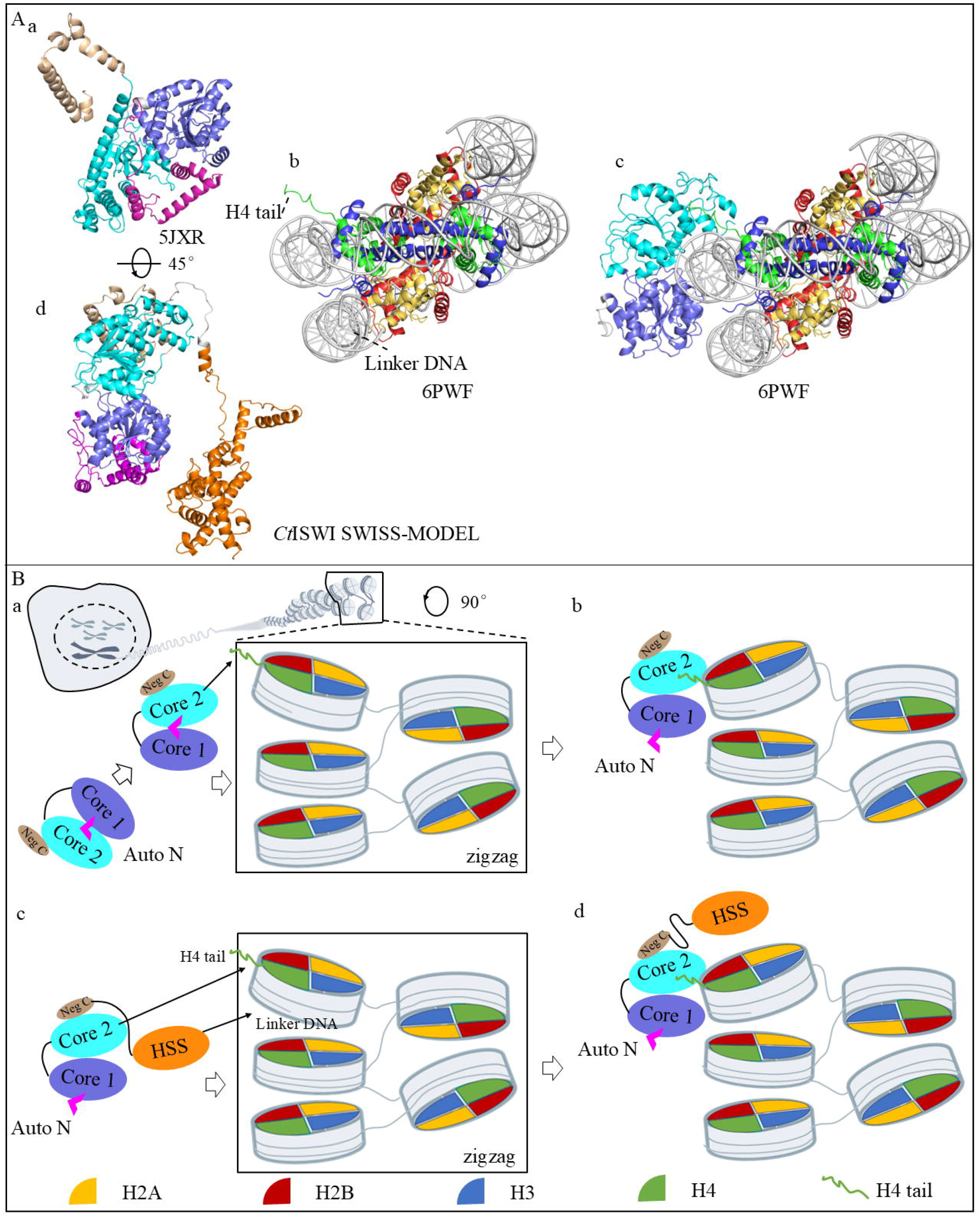
Possible mechanism of stimulating the activity of ISWI. **(A)** Conformational changes of ISWI upon nucleosome binding. **(a)** Crystal structure of MtISWI_81-723_ (PDB: 5JXR). **(b)** Structure of mononucleosome (PDB:6PWF) **(c)** Structure of *Ct*ATPase_77-722_ bound with nucleosome (PDB: 6PWF), where the position of HSS domain is occupied by the nucleosome. **(d)** Structural model 1 of *Ct*ISWI_77-1038_ predicted by SWISS-MODEL. **(B)** Possible conformational changes of ISWI upon binding with nucleosome. **(a)** Conformation of core 1 and core 2 locked together by AutoN. **(b)** Upon binding with nucleosome, the H4 tail inserts into core 2, and AutoN undergoes a significant conformational change. **(c)** ISWI approaches nucleosomes arranged in a “zigzag” pattern. HSS domain that is located above core 1 and core 2, is engaged by linker DNA. **(d)** Core1 and core2 bind to nucleosome DNA and H4 tail binds to core 2, while HSS domain undergoes a conformational change, moving away from the nucleosome.

After ISWI binds to the nucleosome, AutoN releases its constraints on ATPase domain (Figure 2B-C, Figure 6B a), causing a large conformational change then exposing the ATP binding site, and ATPase domain can bind to SHL2 on nucleosome (Figure 6B b). Following the relief of inhibition, the activation of ISWI is completed, then ISWI proceeds to slide the nucleosome using the energy from ATP hydrolysis. If linker DNA is longer than 20 bp, HSS domain will recognize and be drawn by linker DNA. Then the HSS is evicted to other locations and replaced by the ATPase domain, which binds to nucleosomes in a heterogeneous, relaxed “zigzag” chromatin structure (Figure 6B a) ^[31]^. On the other side, H4 tail inserts into the acidic patch of core 2 (Figure 2B-C, Supplementary data Figure S4), finally relieving the auto-inhibition of ISWI (Figure 6B d).

## Supplementary Materials

The following is the Supplementary data to this article.

## Acknowledgements

This work was supported by National Natural Science Foundation of China (NSFC, 31970669) and Research Start Funding of Anhui University (S020318006/067).

## Author Contributions

**Jingjun Hong:** Conceptual and experimental design; supervision; methodology; investigation; data analysis; funding acquisition; writing--review and editing;

**Yiming Zhao:** Methodology; investigation; data collection; visualization; writing–original draft;

**Wansen Tan:** Data analysis; visualization; writing–original draft.

## Data Availability Statement

The data that support the findings of this study are available from the corresponding author upon reasonable request.

