## Supporting Information for "Regulation of CtISWI activity by AutoN and HSS domain"

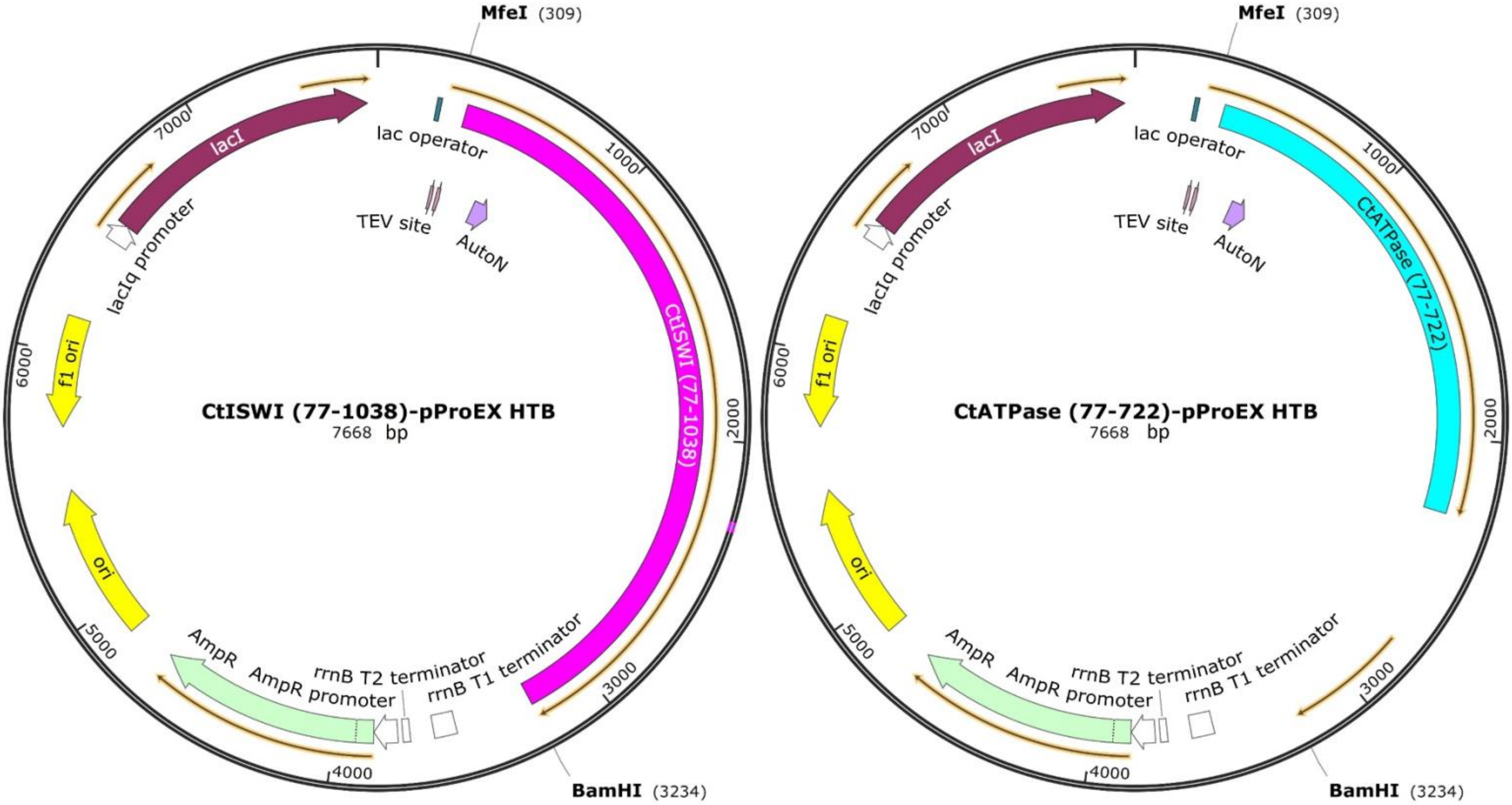


**Figure S1. Plasmid profile analysis of *Ct*ISWI$_{77\text{-}1038}$ and *Ct*ATPase$_{77\text{-}722}$.** pProEx-Htb was used for backbone. *Ct*ISWI$_{77\text{-}1038}$ was inserted between *Mfe* I and *BamH* I. *Ct*ATPase$_{77\text{-}722}$ was constructed through inserting a termination codon before Ala723 of *Ct*ISWI$_{77\text{-}1038}$ to suppress subsequent sequence expression.

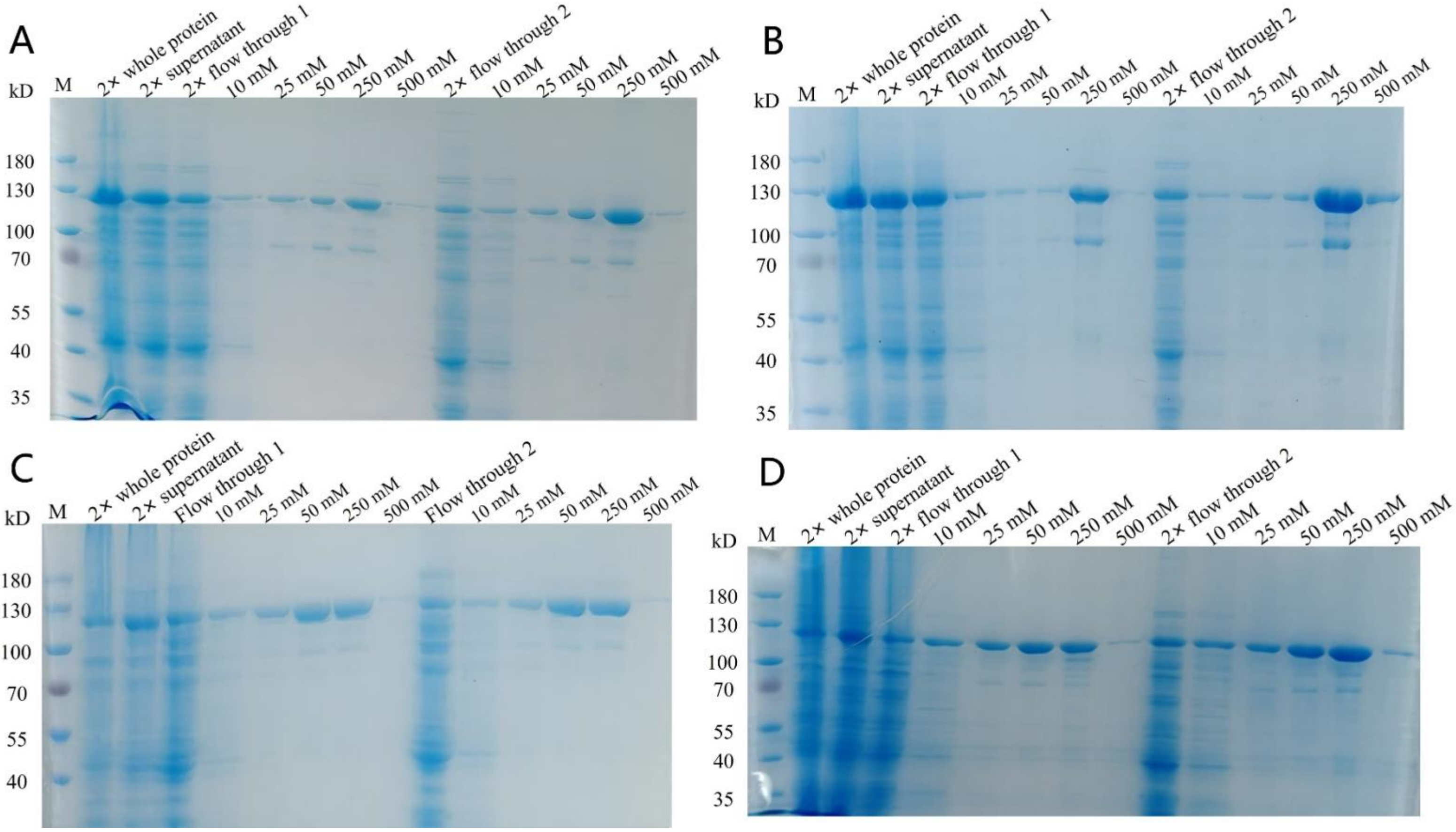


**Figure S2. Ni-affinity chromatography of four *Ct*ISWI mutants.** **(A)** Gels of CtISWI$^{4R/4G}$ purification. **(B)** Gels of CtISWI$^{8G}$ purification. **(C)** Gels of CtISWI$^{8G/4R140A}$ purification. **(D)** Gels of CtISWI$^{8G/3A}$ purification.

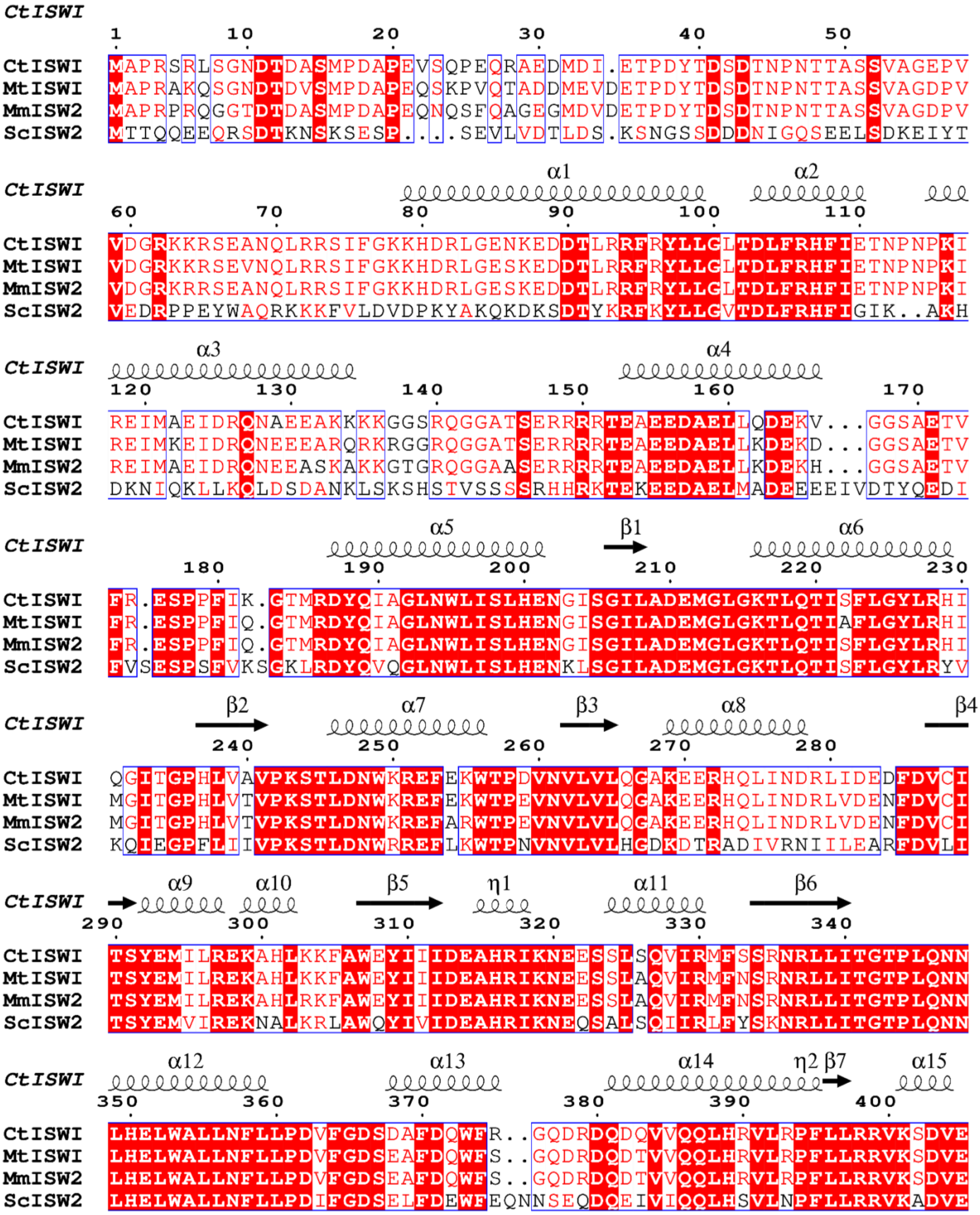
CtISWI
1 10 20 30 40 50
CtISWI MAPRSRLSGNDTDASMPDAPEVSQPEQRAEDMDI.ETPDYTDSDTNPNTTASSVAGEPV
MtISWI MAPRAKQSGNDTDVSMPDAPEQSKPVQTADDMEVDETPDYTDSDTNPNTTASSVAGDPV
MmISW2 MAPRPRQGGTDTDASMPDAPEQNQSFQAGEGMDVDETPDYTDSDTNPNTTASSVAGDPV
ScISW2 MTTQQEEQRSDTKNSKSESP....SEVLVDTLDS.KSNGSSDDDNIGQSEELSDKEIYT
CtISWI α1 α2
60 70 80 90 100 110
CtISWI VDGRKKRSEANQLRRSIFGKKHDRLGENKEDDTLRRFRYLLGLTDLFRHFIETNPNPKI
MtISWI VDGRKKRSEVNQLRRSIFGKKHDRLGESKEDDTLRRFRYLLGLTDLFRHFIETNPNPKI
MmISW2 VDGRKRRSEANQLRRSIFGKKHDRLGESKEDDTLRRFRYLLGLTDLFRHFIETNPNPKI
ScISW2 VEDRPPEYWAQRKKKFVLDVDPKYAKQKDKSDTYKRFKYLLGVTDLFRHFIGIK..AKH
CtISWI α3 α4
120 130 140 150 160 170
CtISWI REIMAEIDRQNAEEAKKKGGSRQGGATSERRRRTEAEEDAELLQDEKV...GGSAETV
MtISWI REIMKEIDRQNEEEARQRKRGGRQGGATSERRRRTEAEEDAELLKDEKD...GGSAETV
MmISW2 REIMAEIDRQNEEASKAKKGTGRQGGAASERRRRTEAEEDAELLKDEKH...GGSAETV
ScISW2 DKNIQKLLKQLDSDANKLSKSHSTVSSSSRHHRKTEKEEDAELMADEEEEIVDTYQEDI
CtISWI α5 β1 α6
180 190 200 210 220 230
CtISWI FR.ESPPFIK.GTMRDYQIAGLNWLISLHENGISGILADEMGLGKTLQTISFLGYLRHI
MtISWI FR.ESPPFIQ.GTMRDYQIAGLNWLISLHENGISGILADEMGLGKTLQTIAFLGYLRHI
MmISW2 FR.ESPPFIQ.GTMRDYQIAGLNWLISLHENGISGILADEMGLGKTLQTISFLGYLRHI
ScISW2 FVSESPSFVKSGKLRDYQVQGLNWLISLHENKLSGILADEMGLGKTLQTISFLGYLRYV
CtISWI β2 α7 β3 α8 β4
240 250 260 270 280
CtISWI QGITGPHLVAVPKSTLDNWKREFEKWTPDVNVLVLQGAKEERHQLINDRLIDEDFDVCI
MtISWI MGITGPHLVTVPKSTLDNWKREFEKWTPEVNVLVLQGAKEERHQLINDRLVDENFDVCI
MmISW2 MGITGPHLVTVPKSTLDNWKREFARWTPEVNVLVLQGAKEERHQLINDRLVDENFDVCI
ScISW2 KQIEGPFLIIVPKSTLDNWRREFLKWTPNVNVLVLHGDKDTRADIVRNIILEARFDVLI
CtISWI α9 α10 β5 η1 α11 β6
290 300 310 320 330 340
CtISWI TSYEMILREKAHLKKFAWEYIIIDEAHRIKNEESSLSQVIRMFSSRNRLLITGTPLQNN
MtISWI TSYEMILREKAHLKKFAWEYIIIDEAHRIKNEESSLAQVIRMFNSRNRLLITGTPLQNN
MmISW2 TSYEMILREKAHLRKFAWEYIIIDEAHRIKNEESSLAQVIRMFNSRNRLLITGTPLQNN
ScISW2 TSYEMVIREKNALKRLAWQYIVIDEAHRIKNEQSALSQIIRLFYSKNRLLITGTPLQNN
CtISWI α12 α13 α14 η2 β7 α15
350 360 370 380 390 400
CtISWI LHELWALLNFLLPDVFGDSDAFDQWFR..GQDRDQDQVVQQLHRVLRPFLLRRVKSDVE
MtISWI LHELWALLNFLLPDVFGDSEAFDQWFS..GQDRDQDTVVQQLHRVLRPFLLRRVKSDVE
MmISW2 LHELWALLNFLLPDVFGDSEAFDQWFS..GQDRDQDTVVQQLHRVLRPFLLRRVKSDVE
ScISW2 LHELWALLNFLLPDIFGDSELFDEWFEQNNSEQDQEIVIQQLHSVLNPFLLRRVKADVE

```
CtISWI   β8 α16 α17 α18
         410 420 430 440 450 460
CtISWI   KSLLPKKEINVYIGMSEMQVKWYKKILEKDIDAVNGAGGKRESKTRLLNIVMQLRKCCN
MtISWI   KSLLPKKEINVYIGMSEMQVKWYQKILEKDIDAVNGAGGKRESKTRLLNIVMQLRKCCN
MmISW2   KSLLPKKEVNVYIGMSEMQVKWYQKILEKDIDAVNGAGGKRESKTRLLNIVMQLRKCCN
ScISW2   KSLLPKIETNVYVGMTDMQIQWYKSLLEKDIDAVNGAVGKREGKTRLLNIVMQLRKCCN

CtISWI   η3 α19 α20 β9 α21
         470 480 490 500 510 520
CtISWI   HPYLFEGAEPGPPYTTDEHLIYNSGKMIVLDKLLKRLQSQGSRVLIFSQMSRLLDILED
MtISWI   HPYLFEGAEPGPPYTTDEHLIYNAGKMVVLDKLLKRIQKQGSRVLIFSQMSRLLDILED
MmISW2   HPYLFEGAEPGPPYTTDEHLVYNSGKMTVLDKLLKRLQKQGSRVLIFSQMSRLLDILED
ScISW2   HPYLFEGAEPGPPYTTDEHLIFNSGKMIILDKLLKRLKEKGSRVLIFSQMSRLLDILED

CtISWI   β10 α22 β11 α23 β12
         530 540 550 560 570 580
CtISWI   YCVFRGYKYCRIDGGTAHEDRIAAIDEYNRPGSDKFIFLLTTRAGGLGINLTTADTVIL
MtISWI   YCVFRGYKYCRIDGSTAHEDRIAAIDEYNKPGSDKFIFLLTTRAGGLGINLTTADIVIL
MmISW2   YCVFRSYKYCRIDGSTAHEDRIAAIDEYNKPDSEKFIFLLTTRAGGLGINLTTADIVIL
ScISW2   YCYFRDFEYCRIDGSTSHEERIEAIDEYNKPNSEKFVFLLTTRAGGLGINLVTADTVIL

CtISWI   α24 β13 α25
         590 600 610 620 630 640
CtISWI   YDSDWNPQADLQAMDRAHRIGQTKQVVVYRFVTDNAIEEKVLERAAQKLRLDQLVIQQG
MtISWI   YDSDWNPQADLQAMDRAHRIGQTKQVVVYRFVTDNAIEEKVLERAAQKLRLDQLVIQQG
MmISW2   YDSDWNPQADLQAMDRAHRIGQTKQVVVYRFVTDNAIEEKVLERAAQKLRLDQLVIQQG
ScISW2   FDSDWNPQADLQAMDRAHRIGQKKQVHVYRFVTENAIEEKVIERAAQKLRLDQLVIQQG

CtISWI   η4 α26 α27 α28 α29
         650 660 670 680 690 700
CtISWI   RAQIATKAAANKEELLSMIQHGAEKVFQTKGAFGLMAEKGANLDDDDIDAILKAGEERT
MtISWI   RAQVAAKAAANKDELLSMIQHGAEKVFQTKGAFGTMAEKGSQLDDDDIDAILQAGETRT
MmISW2   RAQVAAKAAANKDELLSMIQHGAEKVFKTKGAFGALAEKGTDLNDDDIDAILQAGESRT
ScISW2   TGKKTASLGNSKDDLLDMIQFGAKNMFEKK......ASKVT..VDADIDDILKKGEQKT

CtISWI   α30 β14 β15
         710 720 730 740 750
CtISWI   RELNAKYEKLGIDDLQKFT...SESAYEWNGEDFAARKKDIGFTWINPSKRERK..EQI
MtISWI   KELNARYEKLGIDDLQKFT...SESAYEWNGEDFAARKKDIGINWINPAKRERK..EQI
MmISW2   KELNARYEKLGIDDLQKFT...SESAYEWNGEDFAARKKDIGINWINPAKRERK..EQV
ScISW2   QELNAKYQSLGLDDLQKFNGIENQSAYEWNGKSFQKKSNDKVVEWINPSRRERRREQTT

CtISWI   α31 η5 α32
         760 770 780 790 800 810
CtISWI   YSIDKYYKQALHSGGRAADSK...PKAPRAPKQIPVHDYQFYPPRLRDLQDRETAWYRK
MtISWI   YSIDKYYKQTFNAGGRAAEAK...PKAPRAPKQVPVHDYQFYPPRLRELQDRETAYYRK
MmISW2   YSIDKYYKQALHTGGRTAEAK...PKAPRAPKQVPVHDYQFYPPRLRDLQDRETAYYRK
ScISW2   YSVDDYYKEIIGGGSKSASKQTPQPKAPRAPKVIHGQDFQFFPKELDALQEKEQLYFKK
```

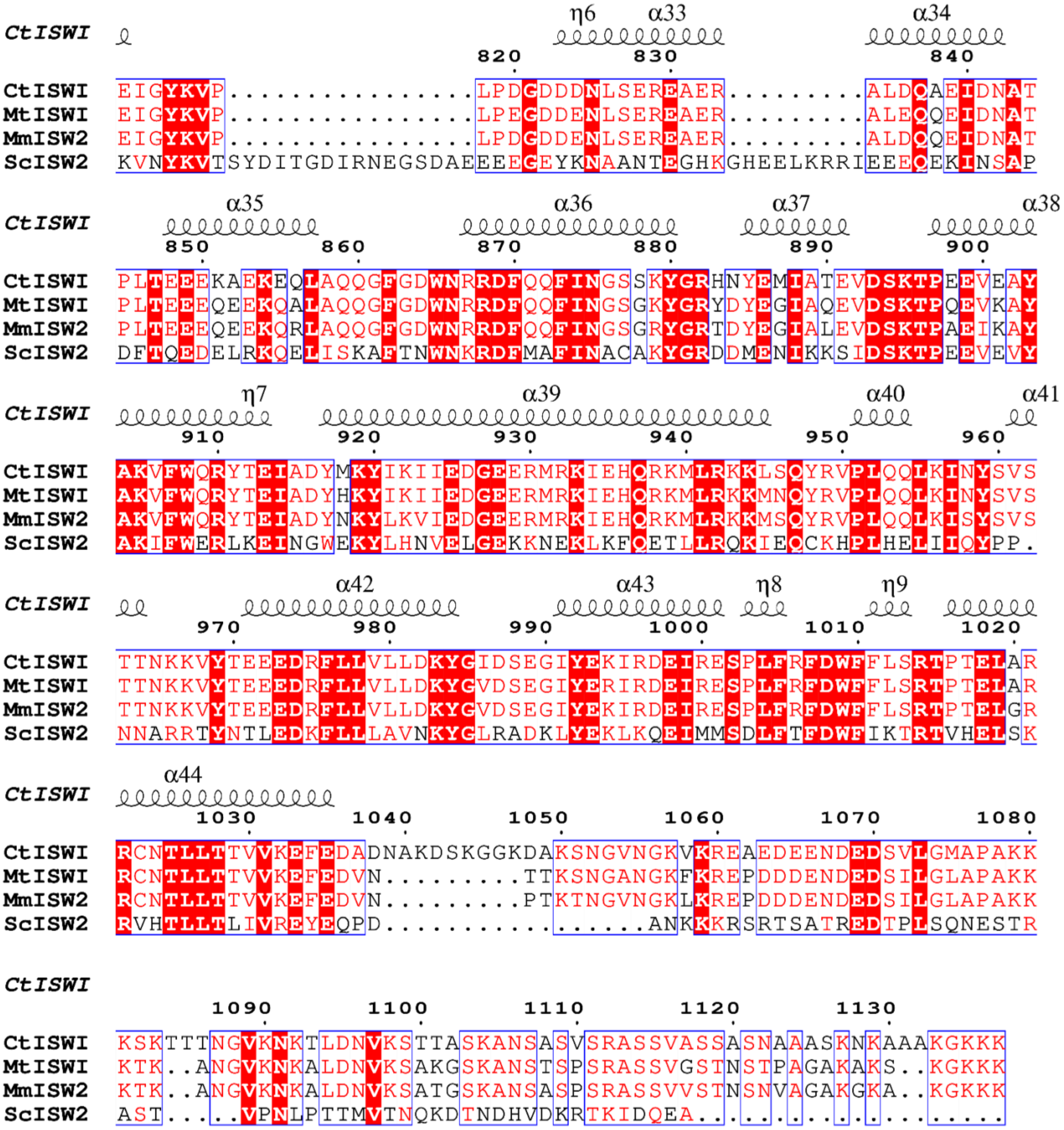

**Figure S3. Sequence alignments of *Ct*ISWI, *Mt*ISWI, *Mm*ISW2, and *Sc*ISW2.** Secondary structural assignments on the top are based on the predicted structural model of our *Ct*ISWI. Strictly conserved residues among all the remodelers included in the alignment are highlighted in white characters on red background.

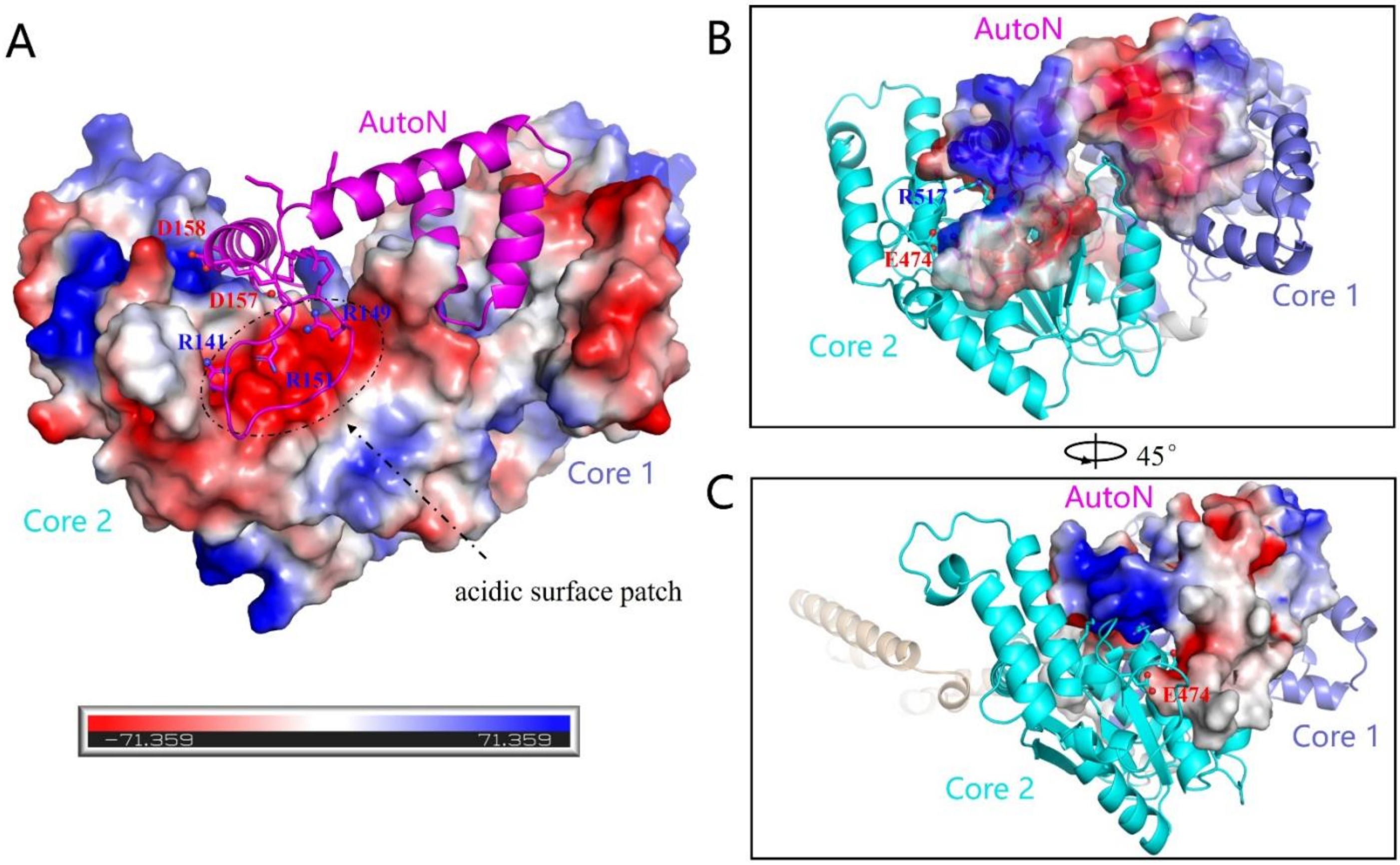


**Figure S4. Mechanism of AutoN binding with core 2. (A)** The AutoN domain is docked onto the core 2 of MtISWI$_{81\text{-}723}$ (PDB: 5JXR) (shown in surface colored using electrostatic potential – blue: positive, red: negative) using coordinates of core 2 as the reference. The acidic patch on core 2 including D520, D524, E523, D537 is marked with circle and arrow. **(B)** Interaction between AutoN and core 2 using coordinates of AutoN as the reference. **(C)** A rotated view to that of the panel (B).

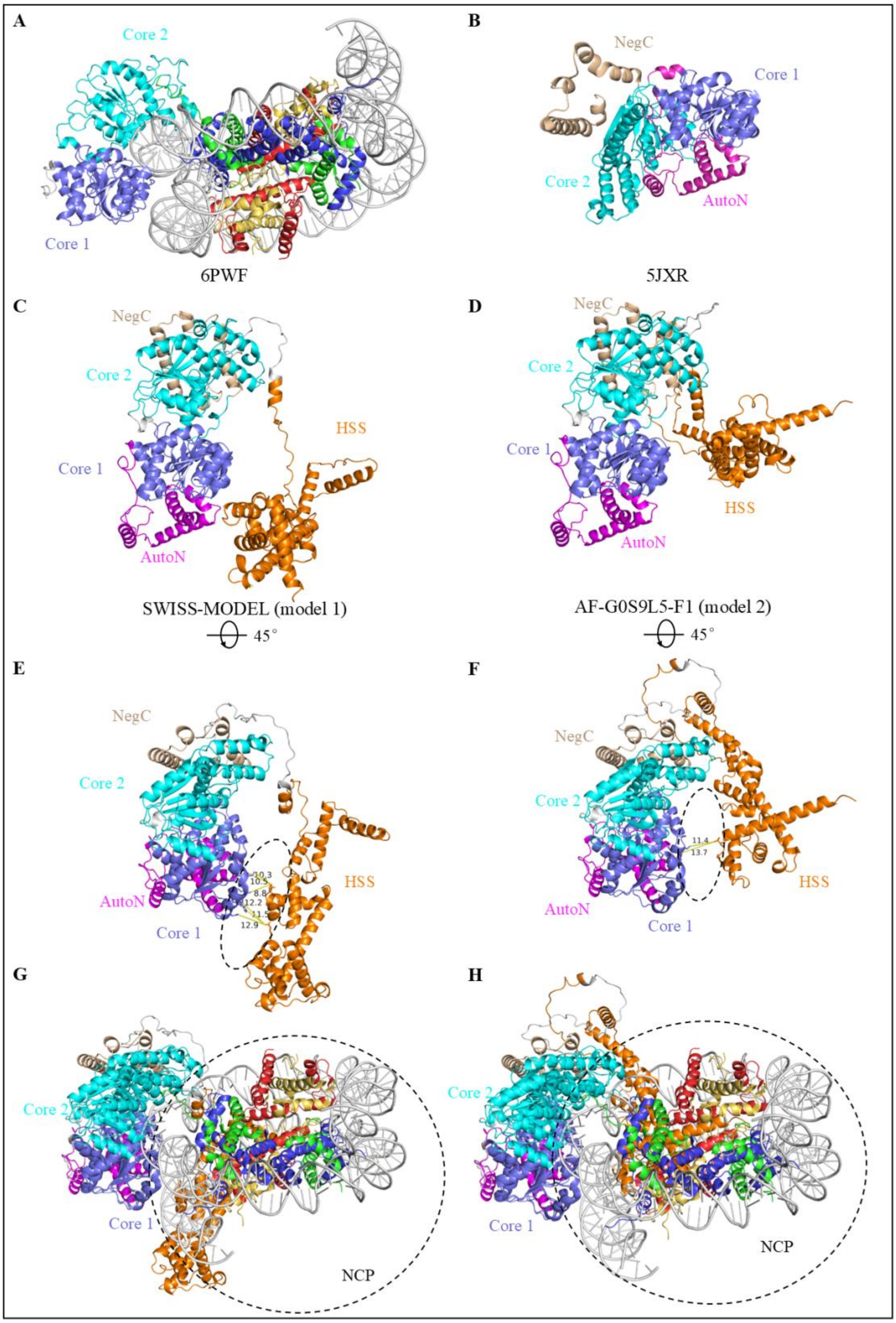
A
Core 2
Core 1
6PWF
B
NegC
Core 1
Core 2
AutoN
5JXR
C
NegC
Core 2
HSS
Core 1
AutoN
SWISS-MODEL (model 1)
45°
D
NegC
Core 2
Core 1
HSS
AutoN
AF-G0S9L5-F1 (model 2)
45°
E
NegC
Core 2
10.3
10.5
8.8
12.2
11.5
12.9
HSS
AutoN
Core 1
F
NegC
Core 2
11.4
13.7
HSS
AutoN
Core 1
G
Core 2
Core 1
NCP
H
Core 2
Core 1
NCP

**Figure S5. Comparison of the cryo-EM structure and predicted model of *Ct*ISWI. (A)** Cryo-EM structure of the nucleosome bound to an ISWI fragment with deletion of AutoN and HSS regions (PDB: 6PWF). (B-D) Structure or model superpositions based on fixing the perspective of core 1. **(B)** X-ray structure of *Mt*ISWI (PDB: 5JXR). **(C)** Structural model of *Ct*ISWI predicted by SWISS-MODEL (model 1). **(D)** Model of *Ct*ISWI predicted by AlphaFold (model 2). **(E)** A rotated view to that of panel (C), and the relatively close distance between HSS and core1 is marked. **(F)** A rotated view to that of panel (D), and the relatively close distance between HSS and core1 is marked. **(G)** Superposition of *Ct*ISWI SWISS-MODEL (E) with 6PWF. Regions where nucleosome conflicts with HSS domain are circled. **(H)** Superposition of AlphaFold model of *Ct*ISWI (F) with 6PWF. Sites where nucleosome conflicts with HSS domain are circled.

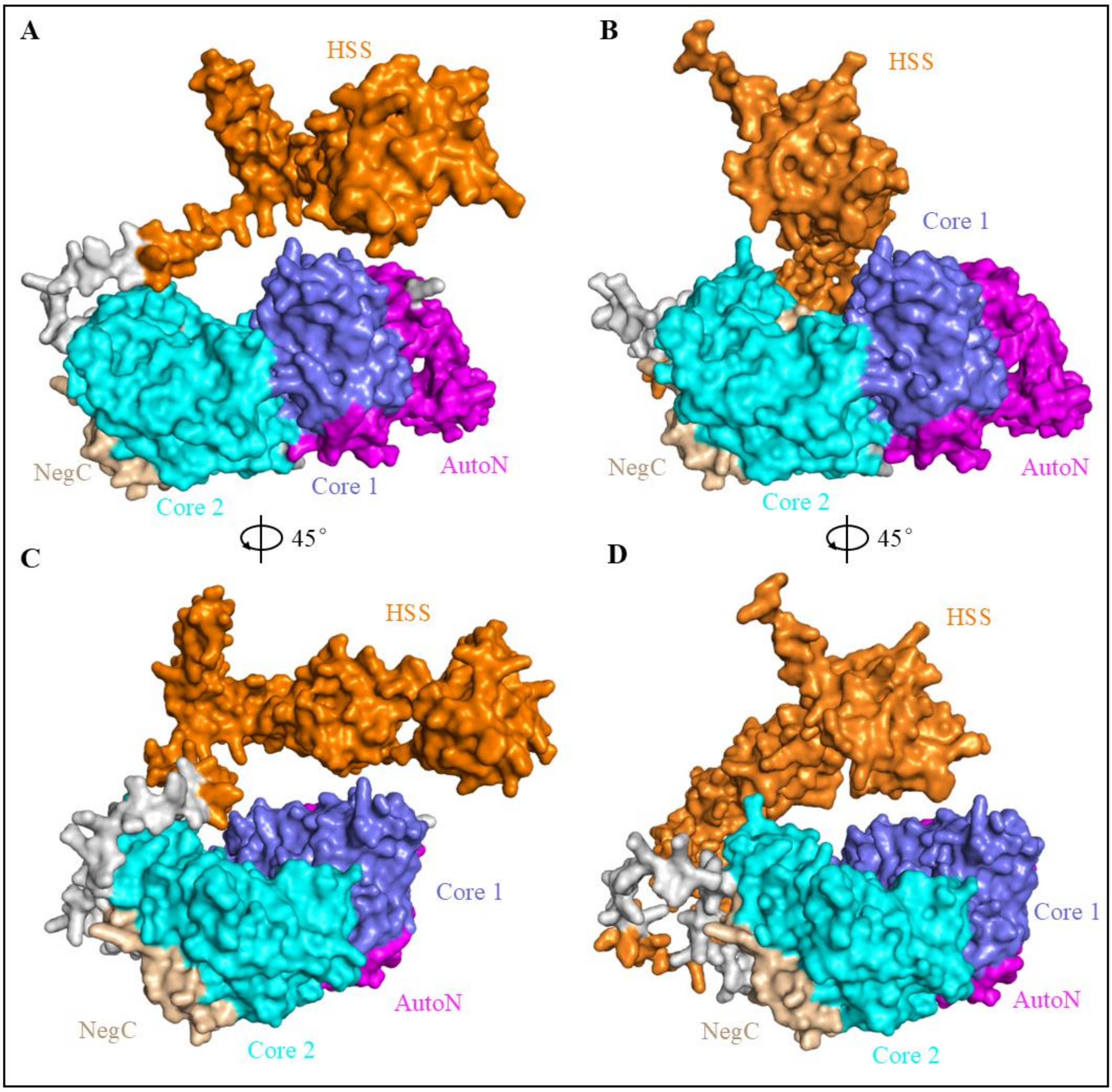


**Figure S6. Surface representation of *CtI*SWI model. (A)** Surface of *Ct*ISWI structure model 1. HSS domain is located closer to core 1 in this model. **(B)** Align Core 1, surface of *Ct*ISWI structure model 2. HSS domain shifts towards core 2 and is located above the junction of core 1 and core 2. **(C)** 45° rotated view to that of panel (A). **(D)** 45° rotated view to that of panel (B). Compared with Core2, the HSS domain is closer to core 1.

**Table S1. PCR procedures for construction of CtISWI$^{4R/4G}$ and CtISWI$^{4K/4G}$.**

| Procedure | Temperature (°C) | Time |
|---|---|---|
| Pre-denaturation | 94 °C | 5 min |
| Denaturation | 95 °C | 60 s |
| Annealing | 55 °C | 60 s |
| Extension | 72 °C | 90 s |
| Cycling | GOTO 2, ×15 | |
| Final extension | 72 °C | 5 min |
| Cooling | 16 °C | Forever |

**Table S2. PCR procedures for construction of CtISWI$^{8G}$.**

| Procedure | Temperature (°C) | Time |
|---|---|---|
| Pre-denaturation | 94 °C | 5 min |
| Denaturation | 95 °C | 60 s |
| Annealing | 60 °C | 60 s |
| Extension | 72 °C | 90 s |
| Cycling | GOTO 2, ×15 | |
| Final extension | 72 °C | 5 min |
| Cooling | 16 °C | Forever |

**Table S3. Two-step PCR procedures for construction of CtISWI$^{8G/R140A}$, CtISWI$^{8G/3A}$ and CtISWI$^{8G/4A}$.**

| Procedure | Temperature (°C) | Time |
|---|---|---|
| Pre-denaturation | 94 °C | 5 min |
| Denaturation | 95 °C | 60 s |
| Annealing and extension | 68 °C | 5 min |
| Cycling | GOTO 2, ×15 | |
| Final extension | 72 °C | 10 min |
| Cooling | 16 °C | Forever |

**Table S4. Primers for protein mutants construction of CtISWI.**

| Primer | Base sequence (5’- 3’) |
|---|---|
| CtISWI 4R/4G (forward) | GGCGCAACCAGTGAAGGTGGCGGTGGCACCGAGGCAGAAGAAGATG |
| CtISWI 4R/4G (reverse) | CATCTTCTTCTGCCTCGGTGCCACCGCCACCTTCACTGGTTGCGCC |
| CtISWI 4K/4G (forward) | CAGAATGCAGAAGAAGCTGGGGGGGGAGGAGGTGGCAGCCGCCAG |
| CtISWI 4K/4G (reverse) | CTGGCGGCTGCCACCTCCTCCCCCCCCAGCTTCTTCTGCATTCTG |
| CtISWI R140/A (forward) | GGAGGAGGTGGCAGCGCCCAGGGTGGCGCAACC |
| CtISWI R140/A (reverse) | GGTTGCGCCACCCTGGGCGCTGCCACCTCCTCC |
| CtISWI E155-D157/A (forward) | GGTGGCACCGAGGCAGCGGCCGCGGCCGAACTGCTGCAA |
| CtISWI E155-D157/A (reverse) | TTGCAGCAGTTCGGCCGCGGCCGCTGCCTCGGTGCCACC |

**Table S5. Primers for dsDNA annealing.**

| Primer | Base sequence (5’- 3’) |
|---|---|
| 18 bp dsDNA (forward) | CAATTGATGGTGATGCCC |
| 18 bp dsDNA (reverse) | AGCAGCAGCGGTAAGAAG |
| 32 bp dsDNA (forward) | GGGCATCACCATCAAGCCAAAATTGAAGAAGG |
| 32 bp dsDNA (reverse) | CTTACCGCTGCTGCTGGCTGCATCGACAGTCTGACG |
| 59 bp dsDNA (forward) | TCCACACTGTTTTACAGCCTTTTCTACTACGTCGTATCAAA ATGGAGGCCCAGAATACC |
| 59 bp dsDNA (reverse) | AGTTCCTTTTTAGGCAGTAAGGATGTTTCCACATCGCT CAGTATAGCGACCAGCATTC |

**Table S6. ssDNA sequence for stimulating ATPase activity.**

| Primer | Base sequence (5'- 3') |
|---|---|
| 33 nt ssDNA | CCAAGGCAGCAGCAGCGACTATCAGATCGCAGG |
| 57 nt ssDNA | CTGCCTGGTGCTGCAAAAAGCGCTGCGAAAGTGGCGGCTAAAGCTCACC ACCACCAC |
| 79 nt ssDNA | GGATAAAATTGTCAAGCAACTCCACACTGTTTTACAGCCTTTTCTACTAC GTCGTATCAAAATGGAGGCCCAGAATACC |

**Table S7. Raw measurement data of ATPase activity of *Ct*ISWI wild-type and several mutant proteins.**

| | WT | 4R/4G | 8G | 8G/R140A | 8G/3A | 8G/4A |
|---|---|---|---|---|---|---|
| 1 | 0.535 | 2.530 | 2.398 | 5.546 | 7.010 | 6.875 |
| 2 | 0.550 | 2.002 | 2.621 | 6.195 | 6.580 | 7.142 |
| 3 | 0.535 | 2.310 | 2.953 | 5.872 | 6.613 | 6.071 |

**Table S8. Raw measurement data of ATPase activity of *Ct*ISWI promoted by dsDNA.**

| | None DNA | +18 bp dsDNA | +32 bp dsDNA | +60 bp dsDNA |
|---|---|---|---|---|
| 1 | 0.156 | 0.137 | 0.431 | 0.763 |
| 2 | 0.110 | 0.254 | 0.489 | 0.842 |
| 3 | 0.0783 | 0.137 | 0.470 | 0.803 |

**Table S9. Raw measurement data of ATPase activity of CtISWI promoted by ssDNA.**

| | None DNA | +33 nt ssDNA | +57 nt ssDNA | +79 nt ssDNA |
|---|---|---|---|---|
| 1 | 0.413 | 2.481 | 1.626 | 1.096 |
| 2 | 0.413 | 2.564 | 1.902 | 1.077 |
| 3 | 0.358 | 2.426 | 1.902 | 1.135 |
